## Supplemental Table 1 for "Vocal complexity in the long calls of Bornean orangutans"

**Table S1.** Table describing features measured in Raven Pro and warbleR (Specan and freq_ts**)**

| **No** | **Program** | **Feature** | **Description** |
| --- | --- | --- | --- |
| 1 | Raven | Delta.Time.s | difference between Begin Time and End Time for the selection (s) |
| 2 | Raven | Freq.5%.Hz | frequency at which summed energy exceeds 5% of total energy |
| 3 | Raven | Freq.95%.Hz | frequency at which summed energy exceeds 95% of total energy |
| 4 | Raven | Agg.Entropy.bits | aggregate entropy measures the disorder in a sound by analyzing the energy distribution (pure tone ~ 0) |
| 5 | Raven | Avg.Entropy.bits | average entropy measurement describes the amount of disorder for a typical spectrum within the selection |
| 6 | Raven | BW.50% | difference between the 25% and 75% frequencies (Hz) |
| 7 | Raven | BW.90% | difference between the 5% and 95% frequencies (Hz) |
| 8 | Raven | Center.Freq | frequency that divides the selection into two frequency intervals of equal energy (Hz) |
| 9 | Raven | Center.Time.Rel. | proportion of selection at which 50% of the sound energy has an earlier time |
| 10 | Raven | Dur.50% | difference between the 25% and 75% times (s) |
| 11 | Raven | Dur.90% | difference between the 5% and 95% times (s) |
| 12 | Raven | Freq.25% | frequency at which summed energy exceeds 25% of total energy (Hz) |
| 13 | Raven | Freq.75% | frequency at which summed energy exceeds 75% of total energy (Hz) |
| 14 | Raven | Peak.Freq | frequency at which Peak Power occurs within the selection (Hz) |
| 15 | Raven | PFC.Avg.Slope | Mean of the Peak Frequency Contour Slope Series of numbers (Hz/ms) |
| 16 | Raven | PFC.Max.Freq | Maximum of the Peak Frequency Contour Series of numbers (Hz) |
| 17 | Raven | PFC.Max.Slope | Maximum of the Peak Frequency Contour Slope Series of numbers (Hz/ms) |
| 18 | Raven | PFC.Min.Freq | Minimum of the Peak Frequency Contour Series of numbers (Hz) |
| 19 | Raven | PFC.Min.Slope | Minimum of the Peak Frequency Contour Slope Series of numbers (Hz/ms) |
| 20 | Raven | PFC.Num.Inf.Pts | Number of times the slope changes sign in Peak Frequency Contour Slope Series of numbers |
| 21 | Raven | Peak.Time.Rel. | proportion of selection at first time in a selection at which amplitude equal to Peak Amplitude occurs |
| 22 | Raven | Time.25%.Rel. | proportion of selection at which 25% of the sound energy has an earlier time |
| 23 | Raven | Time.5%.Rel. | proportion of selection at which 5% of the sound energy has an earlier time |
| 24 | Raven | Time.75%.Rel. | proportion of selection at which 75% of the sound energy has an earlier time |
| 25 | Raven | Time.95%.Rel. | proportion of selection at which 95% of the sound energy has an earlier time |
| 26 | specan | meanfreq | mean of frequency spectrum (kHz) |
| 27 | specan | sd | standard deviation of frequency (kHz) |
| 28 | specan | skew | skewness: asymmetry of the spectrum |
| 29 | specan | kurt | kurtosis: peakedness of the spectrum |
| 30 | specan | sp.ent | energy distribution of the frequency spectrum (pure tone ~ 0) |
| 31 | specan | time.ent | energy distribution on the time envelope (pure tone ~ 0) |
| 32 | specan | entropy | spectrographic entropy: product of time x spectral entropy |
| 33 | specan | sfm | spectral flatness (pure tone ~ 0) |
| 34 | specan | meandom | average of dominant frequency measured across the acoustic signal |
| 35 | specan | mindom | minimum of dominant frequency measured across the acoustic signal |
| 36 | specan | maxdom | maximum of dominant frequency measured across the acoustic signal |
| 37 | specan | dfrange | range of dominant frequency measured across the acoustic signal |
| 38 | specan | modindx | modulation index: cumulative difference between adjacent dominant frequencies / dominant frequency range |
| 39 | specan | startdom | dominant frequency measurement at the start of the signal |
| 40 | specan | enddom | dominant frequency measurement at the end of the signal |
| 41 | specan | dfslope | slope of the change in dominant frequency through time |
| 42 | specan | meanpeakf | frequency with highest energy from the mean frequency spectrum |
| 43 | specan | Freq_IQR | interquartile frequency range. Frequency range between 'freq.Q25' and 'freq.Q75' (kHz) |
| 44 | specan | Time_IQR | interquartile time range. Time range between 'time.Q25' and 'time.Q75' (s) |
| 45 | freq_ts | F0_min | frequency at which F0 contour is at its minimum value (kHz) |
| 46 | freq_ts | F0_max | frequency at which F0 contour reaches its maximum value (kHz) |
