## Supplemental Table 2a for "Vocal complexity in the long calls of Bornean orangutans"

**Table S2a.** Table summarizing results of Kruskal-Wallis tests for differences among pulses or clusters identified by human observers, affinity propagation, and fuzzy clustering for each of the top five influential variables.

|  | **A/V** | | | **AFFINITY** | | | **FUZZY** | | |
| --- | --- | --- | --- | --- | --- | --- | --- | --- | --- |
| **Variable** | **χ²** | **df** | **p** | **χ²** | **df** | **p** | **χ²** | **df** | **p** |
| Center | 557.81 | 5.00 | 0.00 | 738.53 | 3.00 | 0.00 | 417.16 | 1.00 | 0.00 |
| Peak | 425.31 | 5.00 | 0.00 | 588.78 | 3.00 | 0.00 | 406.49 | 1.00 | 0.00 |
| Mean peak | 528.18 | 5.00 | 0.00 | 677.78 | 3.00 | 0.00 | 421.95 | 1.00 | 0.00 |
| Third quart | 570.30 | 5.00 | 0.00 | 777.79 | 3.00 | 0.00 | 416.69 | 1.00 | 0.00 |
| First quart | 536.72 | 5.00 | 0.00 | 684.50 | 3.00 | 0.00 | 414.47 | 1.00 | 0.00 |
