## Supplemental Table 2b for "Vocal complexity in the long calls of Bornean orangutans"

**Table S2b.** Table summarizing results of Dunn tests for pair-wise differences among pulses identified by human observers for each of the top five influential variables.

| **A/V** | **Center** | | | **Peak** | | | **Mean peak** | | | **Third quart** | | | **First quart** | | |
| --- | --- | --- | --- | --- | --- | --- | --- | --- | --- | --- | --- | --- | --- | --- | --- |
| Pair | Z | P.unadj | P.adj | Z | P.unadj | P.adj | Z | P.unadj | P.adj | Z | P.unadj | P.adj | Z | P.unadj | P.adj |
| HR-HU | -0.21 | 0.84 | 0.84 | -0.54 | 0.59 | 0.63 | 0.12 | 0.90 | 0.90 | -0.60 | 0.55 | 0.64 | -0.54 | 0.59 | 0.68 |
| HR-IN | 10.29 | 0.00 | 0.00 | 9.12 | 0.00 | 0.00 | 10.59 | 0.00 | 0.00 | 10.52 | 0.00 | 0.00 | 9.04 | 0.00 | 0.00 |
| HR-LR | 9.80 | 0.00 | 0.00 | 9.81 | 0.00 | 0.00 | 9.68 | 0.00 | 0.00 | 9.97 | 0.00 | 0.00 | 9.35 | 0.00 | 0.00 |
| HR-SI | 19.72 | 0.00 | 0.00 | 17.10 | 0.00 | 0.00 | 19.35 | 0.00 | 0.00 | 19.86 | 0.00 | 0.00 | 19.26 | 0.00 | 0.00 |
| HR-VO | -0.67 | 0.50 | 0.58 | -0.84 | 0.40 | 0.50 | -0.44 | 0.66 | 0.71 | -0.55 | 0.58 | 0.62 | -0.50 | 0.62 | 0.66 |
| HU-IN | 7.12 | 0.00 | 0.00 | 6.65 | 0.00 | 0.00 | 7.00 | 0.00 | 0.00 | 7.65 | 0.00 | 0.00 | 6.60 | 0.00 | 0.00 |
| HU-LR | 6.31 | 0.00 | 0.00 | 6.66 | 0.00 | 0.00 | 5.90 | 0.00 | 0.00 | 6.81 | 0.00 | 0.00 | 6.37 | 0.00 | 0.00 |
| HU-SI | 11.40 | 0.00 | 0.00 | 10.27 | 0.00 | 0.00 | 10.83 | 0.00 | 0.00 | 11.90 | 0.00 | 0.00 | 11.49 | 0.00 | 0.00 |
| HU-VO | -0.36 | 0.72 | 0.77 | -0.23 | 0.82 | 0.82 | -0.44 | 0.66 | 0.76 | 0.04 | 0.97 | 0.97 | 0.04 | 0.97 | 0.97 |
| IN-LR | -1.83 | 0.07 | 0.08 | -0.57 | 0.57 | 0.66 | -2.26 | 0.02 | 0.03 | -1.91 | 0.06 | 0.07 | -0.91 | 0.36 | 0.45 |
| IN-SI | 5.60 | 0.00 | 0.00 | 4.63 | 0.00 | 0.00 | 4.91 | 0.00 | 0.00 | 5.46 | 0.00 | 0.00 | 6.70 | 0.00 | 0.00 |
| IN-VO | -7.70 | 0.00 | 0.00 | -7.06 | 0.00 | 0.00 | -7.67 | 0.00 | 0.00 | -7.74 | 0.00 | 0.00 | -6.67 | 0.00 | 0.00 |
| LR-SI | 9.44 | 0.00 | 0.00 | 6.48 | 0.00 | 0.00 | 9.18 | 0.00 | 0.00 | 9.38 | 0.00 | 0.00 | 9.51 | 0.00 | 0.00 |
| LR-VO | -6.92 | 0.00 | 0.00 | -7.09 | 0.00 | 0.00 | -6.60 | 0.00 | 0.00 | -6.90 | 0.00 | 0.00 | -6.45 | 0.00 | 0.00 |
| SI-VO | -12.16 | 0.00 | 0.00 | -10.83 | 0.00 | 0.00 | -11.70 | 0.00 | 0.00 | -12.12 | 0.00 | 0.00 | -11.71 | 0.00 | 0.00 |
