## Supplemental Table 2c for "Vocal complexity in the long calls of Bornean orangutans"

**Table S2c.** Table summarizing results of Dunn tests for pair-wise differences among clusters identified by affinity propagation for each of the top five influential variables.

| **AFFINITY** | **Center** | | | **Peak** | | | **Mean peak** | | | **Third quart** | | | **First quart** | | |
| --- | --- | --- | --- | --- | --- | --- | --- | --- | --- | --- | --- | --- | --- | --- | --- |
| Pair | Z | P.unadj | P.adj | Z | P.unadj | P.adj | Z | P.unadj | P.adj | Z | P.unadj | P.adj | Z | P.unadj | P.adj |
| 152-16 | 0.84 | 0.40 | 0.40 | -0.43 | 0.66 | 0.66 | 0.24 | 0.81 | 0.81 | 2.33 | 0.02 | 0.02 | 0.20 | 0.84 | 0.84 |
| 152-616 | 16.64 | 0.00 | 0.00 | 14.23 | 0.00 | 0.00 | 15.60 | 0.00 | 0.00 | 18.09 | 0.00 | 0.00 | 15.68 | 0.00 | 0.00 |
| 16-616 | 22.88 | 0.00 | 0.00 | 21.43 | 0.00 | 0.00 | 22.33 | 0.00 | 0.00 | 22.56 | 0.00 | 0.00 | 22.51 | 0.00 | 0.00 |
| 152-812 | 7.95 | 0.00 | 0.00 | 7.59 | 0.00 | 0.00 | 7.57 | 0.00 | 0.00 | 8.88 | 0.00 | 0.00 | 7.73 | 0.00 | 0.00 |
| 16-812 | 9.94 | 0.00 | 0.00 | 11.37 | 0.00 | 0.00 | 10.31 | 0.00 | 0.00 | 8.97 | 0.00 | 0.00 | 10.60 | 0.00 | 0.00 |
| 616-812 | -15.61 | 0.00 | 0.00 | -11.83 | 0.00 | 0.00 | -14.41 | 0.00 | 0.00 | -16.52 | 0.00 | 0.00 | -14.26 | 0.00 | 0.00 |
