## Supplementary figures and images for "Vocal complexity in the long calls of Bornean orangutans"

### Supplemental Figure 1

**Figure S1.** Example of annotated spectrogram

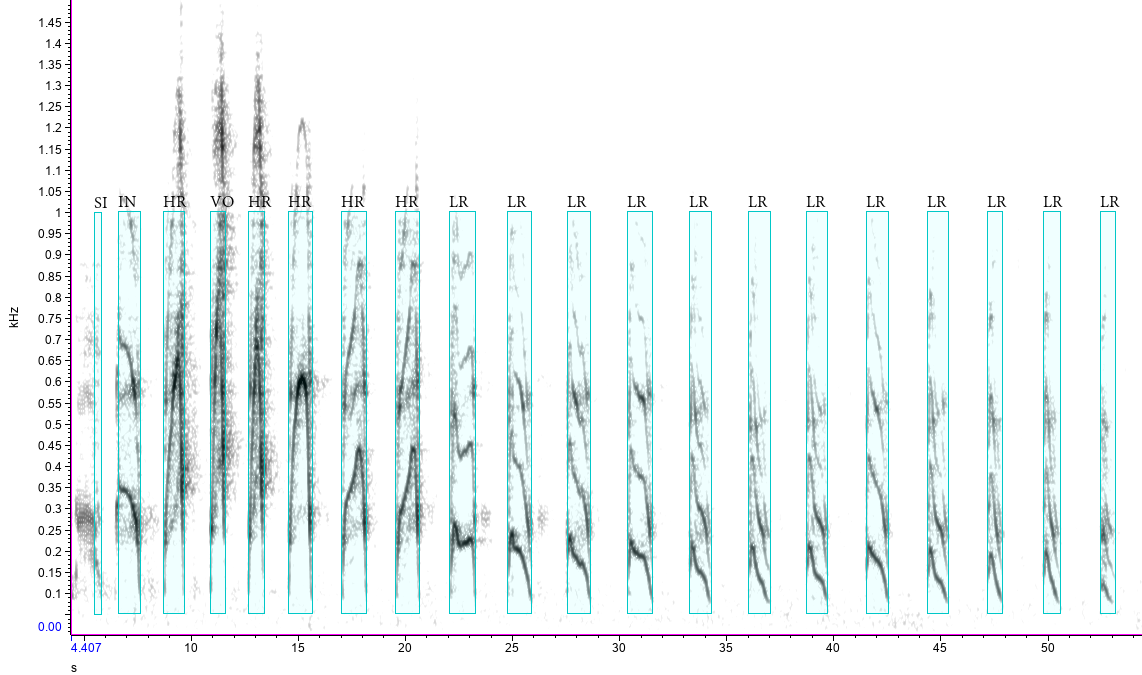
