## Supplemental Figure 2 for "Vocal complexity in the long calls of Bornean orangutans"

**Figure S2.** Boxplots of features that differed across human-labeled pulses (upper left), affinity propagation clusters (upper right), and typical calls in fuzzy clusters (lower left) for each of the following influential features: a) center frequency, b) peak frequency, c) mean peak frequency, d) third quartile frequency, e) first quartile frequency.

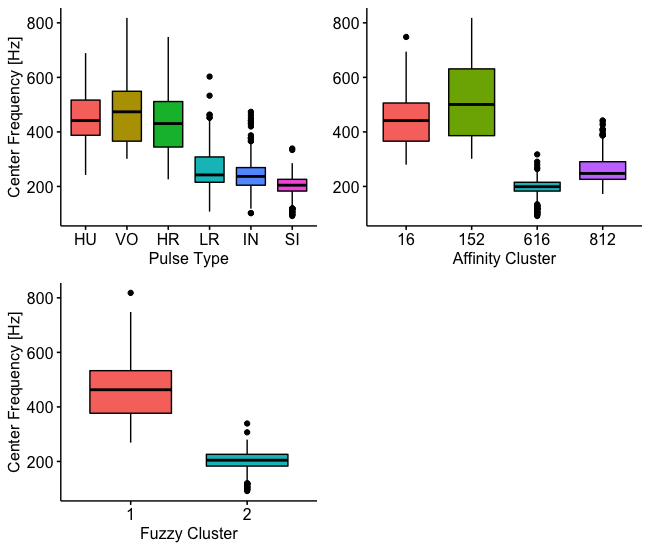

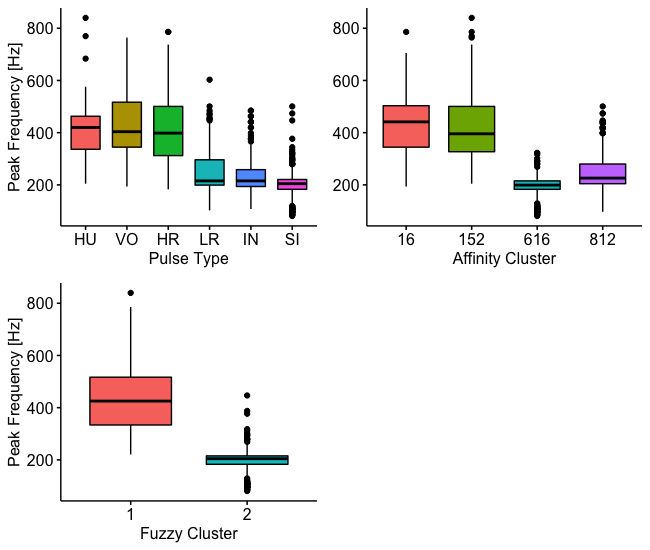

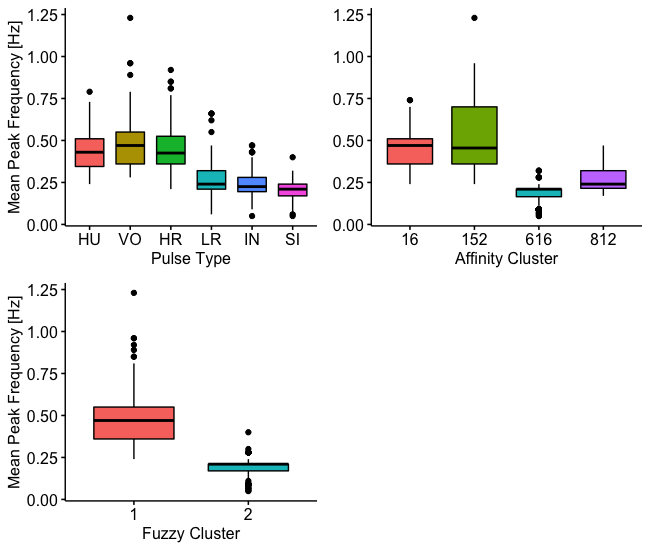

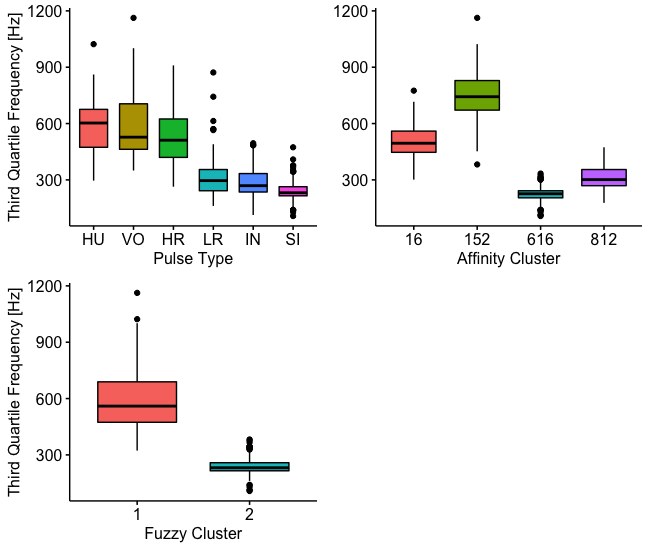

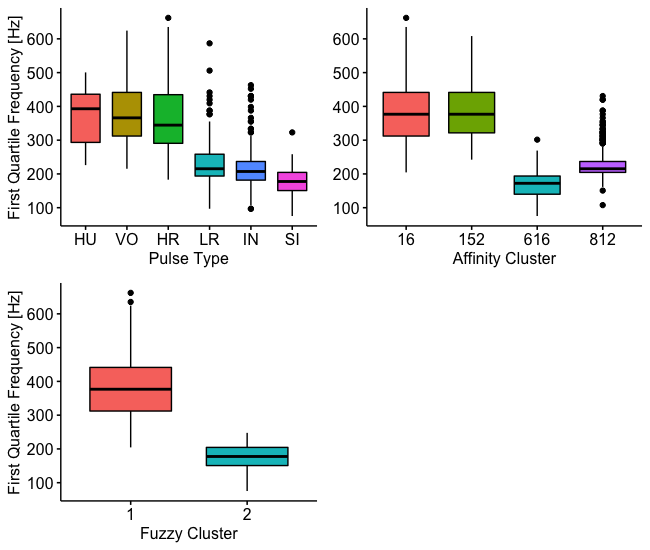
