## Supplemental Figure 3 for "Vocal complexity in the long calls of Bornean orangutans"

**Figure S3.** Histograms (a-b) and error plots (c-d) showing bootstrapping results across 25 iterations within 100–900 randomly sampled observations showing: a) distribution of the number of clusters identified by affinity propagation, b) distribution of the number of clusters identified by fuzzy clustering, c) mean and standard error of typicality coefficients from fuzzy clustering solutions, d) mean and standard error of the classification accuracy from support vector machines.

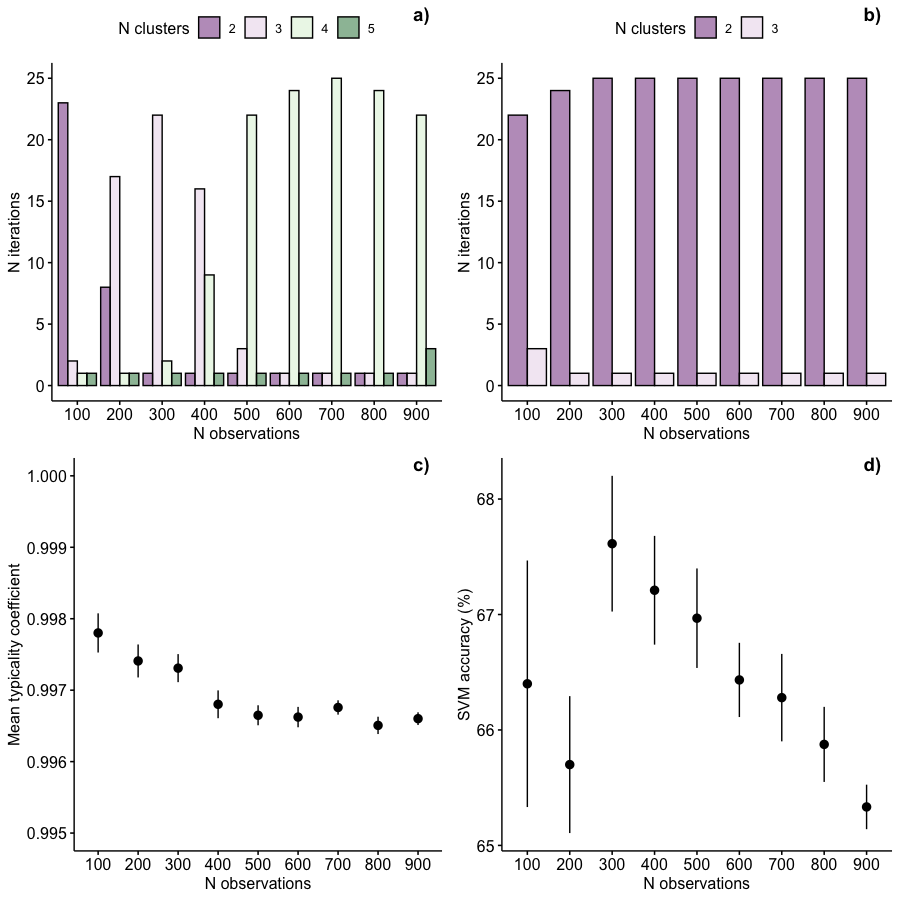
