## Supplemental Figure 4 for "Vocal complexity in the long calls of Bornean orangutans"

**Figure S4.** Histograms showing bootstrapping results across 25 iterations within 2–40 randomly sampled features showing: a) distribution of the number of clusters identified by affinity propagation and b) distribution of the number of clusters identified by fuzzy clustering.

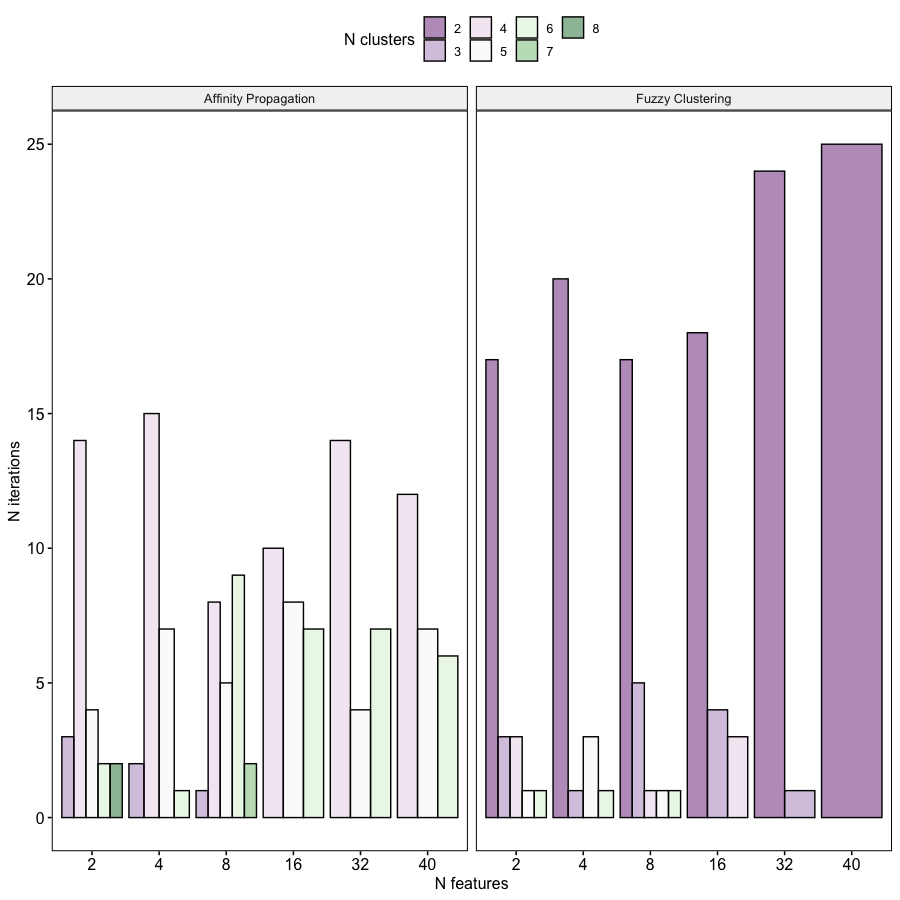
